# IcmX Plug Ejection in *L. pneumophila*’s Type IV Secretion System: Is It Possible?

**DOI:** 10.64898/2025.11.30.691423

**Authors:** Cayson Hamilton, Samuel Bills, Brenden Stark, Gus L.W. Hart

**Affiliations:** Brigham Young University; Georgia Institute of Technology

## Abstract

*Legionella pneumophila* is a gram-negative bacterial pathogen that is the causative agent of several infectious diseases, notably the severe form of pneumonia known as Legionnaires’ Disease. *L. pneumophila* targets amoebas and, in humans, alveolar macrophages. *L. pneumophila* infects its host by envelopment into the host cell through phagocytosis followed by the activation of the Type IV Secretion System (T4SS). The T4SS secretes effector proteins into the host cell which deactivate cell defenses and reprogram cell function to support *L. pneumophila* reproduction. After reproduction, *L. pneumophila* lyses the host cell and the cycle repeats, causing swelling and destroying a primary cell in the host’s immune system—infections caused by *L. pneumophila* are notoriously hard to treat. The stages of the infection process are known but the physical mechanisms are poorly understood. The structure of the T4SS includes a 13 member polymer, DotG which protrudes from the outer leaflet of the bacterial outer membrane. Given this exposure and the necessity of T4SS for infection, DotG is a promising drug target. However, designing an effective drug requires understanding the physical mechanisms. Via MD simulations we have tested hypotheses regarding its function. Specifically, we show the feasibility that an applied force on IcmX, a plug-like penta-mer that initially blocks transport through the T4SS channel, can cause a structural change in DotG and allow for protein release. Further, this applied force could reasonably come from a build up of pressure in the channel from other effector proteins. We conclude that the process of secretion is dependent on the induced structural change in DotG. Inhibiting this change provides a possible drug development direction.

## 2 Introduction

*Legionella pneumophila* are gram-negative opportunistic pathogens that manipulate host cell processes to recruit nutrients and replicate. They are most often found in water sources both, including lakes, reservoirs, fountains, pools, or even air conditioning units [1]. In humans, *L. pneumophila* primarily targets alveolar macrophages, inducing a severe and potentially fatal form of pneumonia known as Legionnaires’ disease [2]. Much is known about *L. pneumophila’s* infection process, which begins with absorption of the pathogen into macrophages through phagocytosis. This absorption is followed by the secretion of over 300 effector proteins which manipulate host cell function. During this process, a protective membrane is formed around the bacteria, known as the *Legionella* Containing Vacuole (LCV), which provides an environment suitable for replication [3, 1]. Studies suggest *L. pneumophila* enters a vegetative, replicative form. After replication, a mature, infectious form is regained, the protective membrane is broken and the host cell is lysed, causing severe swelling and tissue damage. The new *L. pneumophila* will then repeat this process [1, 4]. These infections are very hard to treat once they have begun, which attributes to Legionnaires’ high mortality rate between 4—18% for those who were previously healthy, and much higher for those who are immunocompromised or have preexisting lung conditions such as asthma or smoking [5].

The Type IV Secretion System is central to the infection process. It is a complex molecular machine which is responsible for the secretion of effector proteins to manipulate host cell behavior. Following envelopment into the host cell, the T4SS establishes distinct contact sites with the LCV. The connection of the T4SS to the LCV membrane creates the channel which enables transportation of the effector proteins into the host cell, as diagrammed in Figure 1 [6]. Targeted gene knock-out studies have shown that when the T4SS is rendered non-functional, the LCV is unable to form and effector proteins are unable to be released into the host cell [6]. A non-functional T4SS renders *L. pneumophila* incapable of inducing any form of infection; they will be eliminated by the alveolar macrophages.

**Figure 1:**
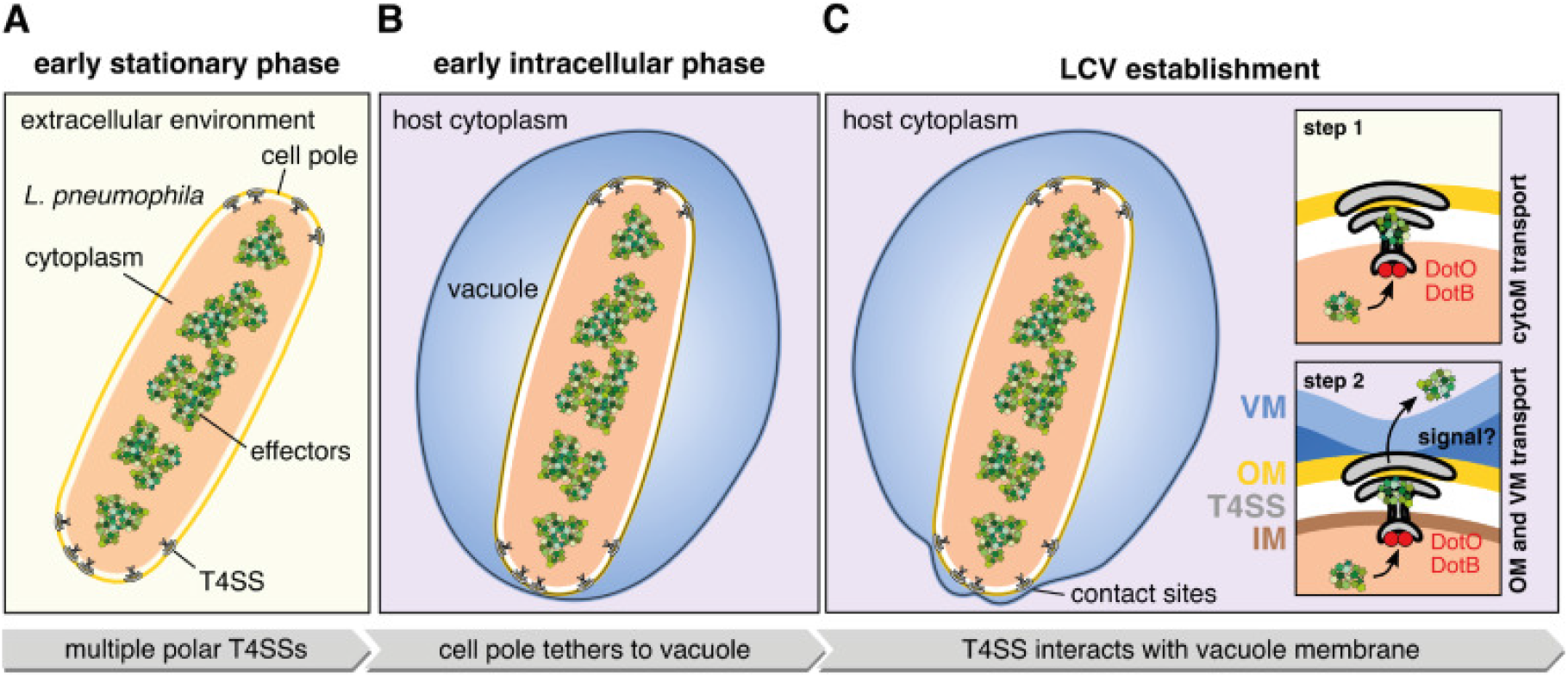
T4SS and LCV Formation: (A) Diagram highlighting the location of the T4SSs in the polar regions of the cell membrane. (B) Once the bacteria has entered the host cell, the cell poles tether to the membrane of the LCV. (C) The T4SS interacts with the vacuole membrane, establishing a site through which effector proteins can be secreted into the host cell. Figure adapted from [6].

The T4SS spans from the bacerial cytoplasm through the inner and outer membranes, with a certain 13-member polymer (DotG) protruding into the host cell cytoplasm (shown in yellow in Figure 2 Panel A). The convenient location of the protein system, combined with its essential nature in inducing infection makes DotG a promising drug target. While much has been discovered in recent years regarding the steps through which infection occurs, little is understood regarding the physical mechanics. Specifically, it is unknown how the proteins make their way out of the T4SS into the host cell. Greater mechanical insight into this process would provide invaluable direction in designing an inhibitory drug.

**Figure 2:**
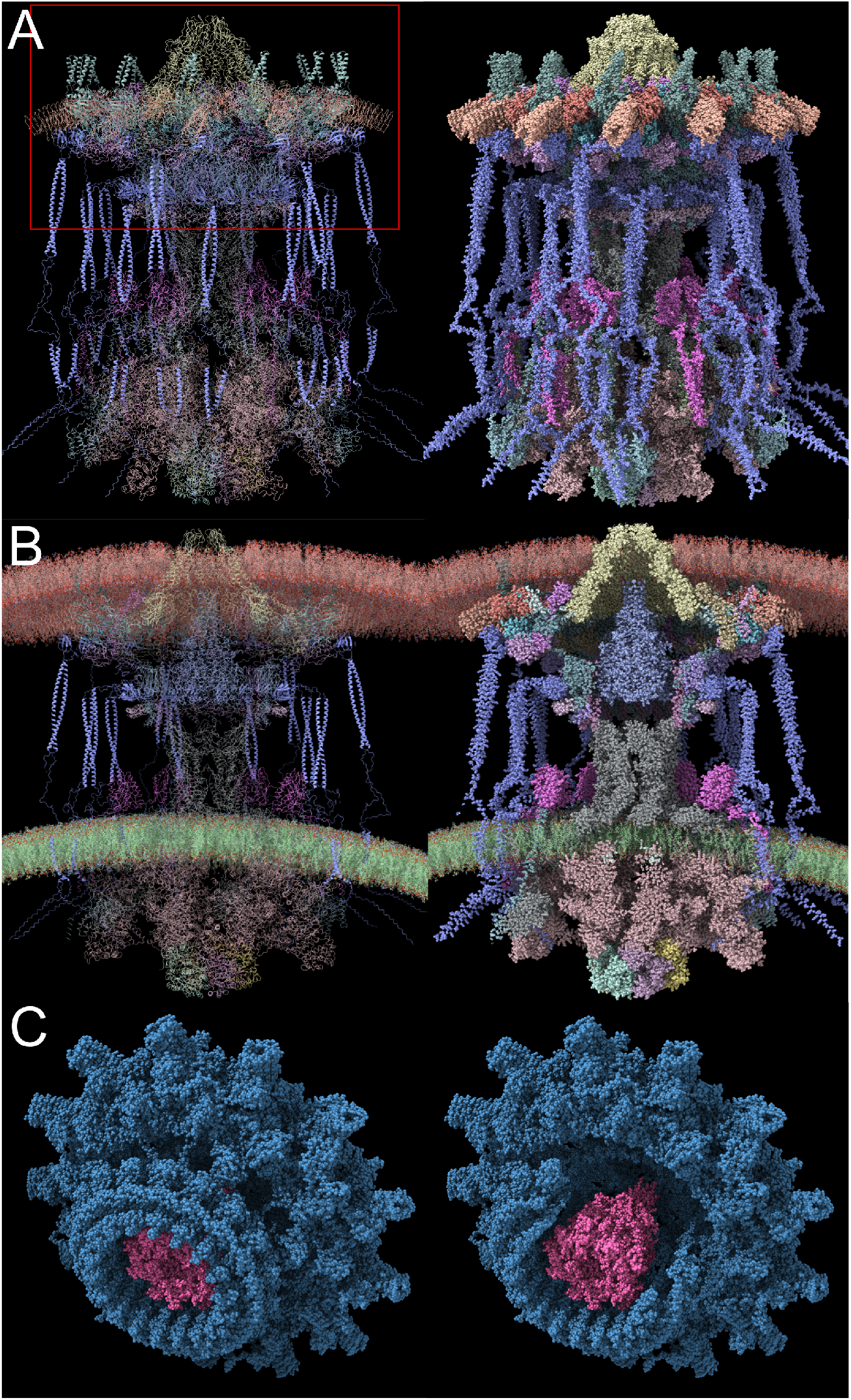
Computational T4SS Model: (A) The T4SS complex – visualizations showing the secondary structures (left) as well as the atomistic view (right). The red rectangle denotes the section in our simulations. (B) A cross section look at the complex, with membranes included for reference. (C) A closer look at the section of interest, particularly highlighting the plug protein (IcmX) in pink. A 360-degree video rotation of the entire complex can be seen here.

Cryo-electron microscopy and tomography combined with gene knock-out testing, have revealed the structure of the T4SS. These recent developments, combined with previous work which to solve the outer membrane complex (OMC, shown in blue in Figure 2 Panel B) [2, 7], led to the completion of a near-complete atomic model of the T4SS [8]. This new model enables further exploration and testing of hypotheses regarding the machine’s function.

Many homologs are apparent with other type IV secretion systems. Specifically, there is a protein, “the plug” (also known as IcmX, shown in pink in Figure 2 Panel B) that resembles the stalk protein in the R388 conjugative T4SS [9]. Figure 3 shows a side-by-side comparison of the two proteins, with homologous proteins indicated. Each of these proteins sit in the secretion channel and appear to block the transport of proteins until they have been removed. The stalk becomes the tip of a pilus that is built by the R388 machine and pushed out through the channel. However, no evidence suggesting that a pilus is created by *L. pneumophila’s* T4SS has ever been found [6]. Another distinct difference between *L. pneumophila’s* T4SS and the R388 T4SS is the extended, mushroom-like OMC in *L. pneumophila*. The added complexity suggests some functionality beyond that of a simple channel. Several hypotheses have been suggested regarding this functionality [8]. Perhaps the OMC might change conformation to contribute to the LCV membrane interaction, facilitating the transportation of effector proteins through the LCV, or perharps the effector proteins may enter the T4SS channel unfolded and the OMC assists with the protein folding. Or, the extended OMC might only provide a structural benefit for the system.

**Figure 3:**
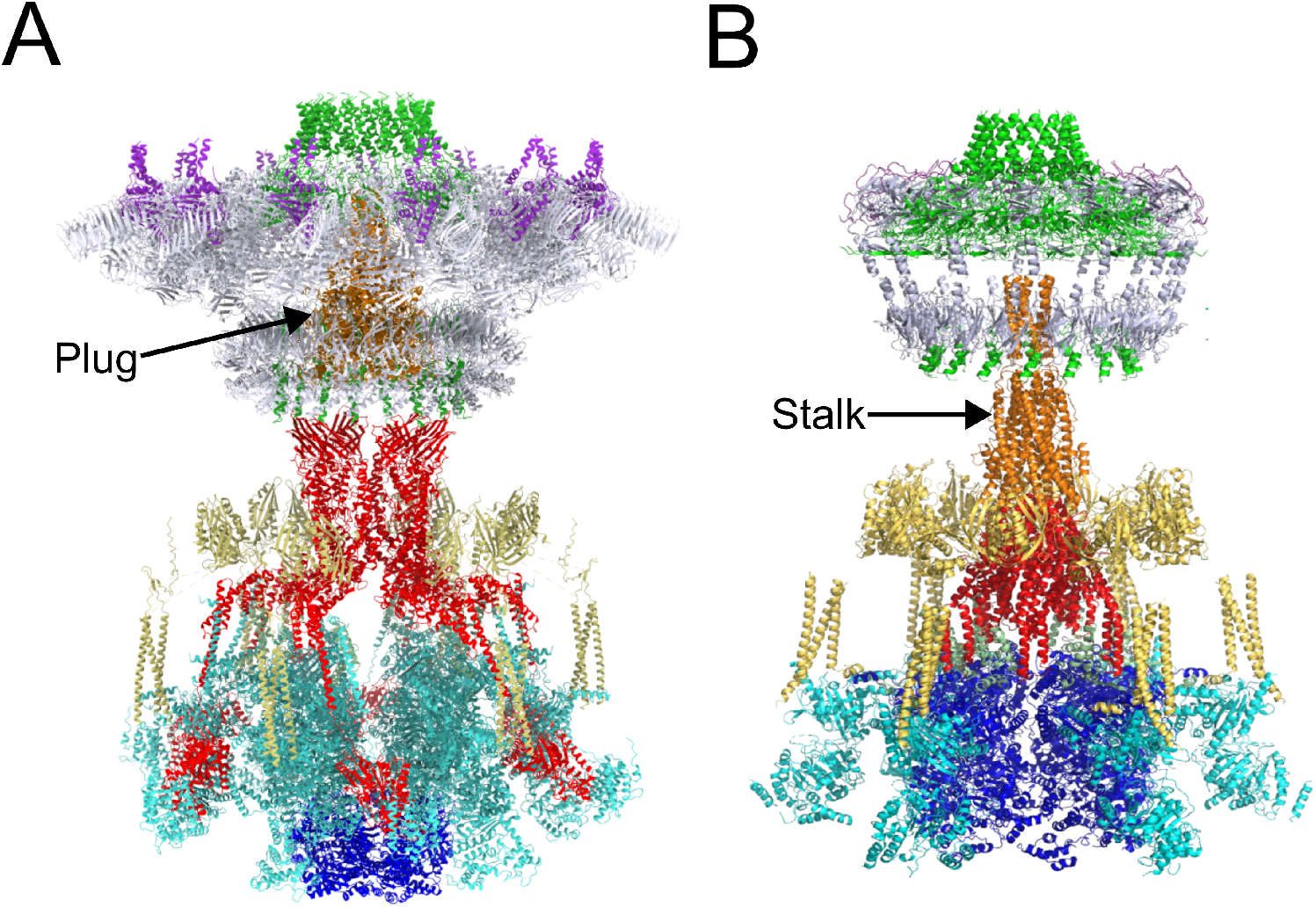
Comparison of IcmX and the R388 Stalk: (A) The *Legionella* Dot/Icm T4SS is shown, with the “plug” protein highlighted in orange. (B) The R388 T4SS is shown, with the “stalk” highlighted in orange, adapted from [9].

**Figure 4:**
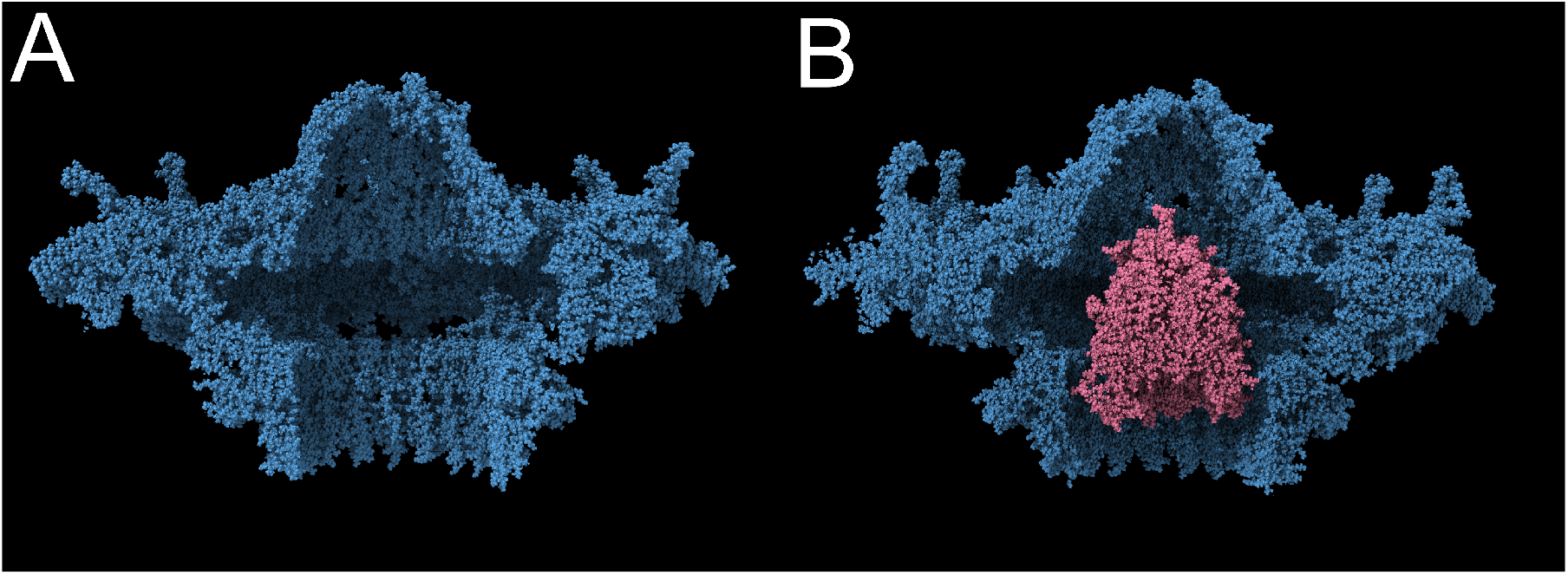
System Structural Comparison. (A) Simulation run without the plug, at 3400 ps. (B) Simulation run with the plug, at 3400 ps.

Whatever the function of the OMC proteins, it is clear that the plug protein must be expelled before effector proteins can be secreted. The plug cannot be removed until *L. pneumophila* is in the correct position as the plug also blocks the entry of foreign molecules. There are, as with the OMC proteins, numerous proposed hypotheses regarding how the plug protein is removed. One builds on the function of the homologous R388 T4SS, which uses its stalk protein as the tip of a pilus [9]. While no pilus is created by *L. pneumophila’s* T4SS, it is possible that an applied force from below pushes the plug protein through the top opening of the OMC, and into the host cell.

Such a force would likely result from effector proteins ready to be secreted and pushed into the channel by the ATPases located on the cytoplasmic section of the T4SS (DotB, DotO) [10]. A structural change in the OMC is necessary for any protein to make its way out of the cell. This is the hypothesis that we sought to test, that if the plug was pushed towards DotG by an external force, DotG would sufficiently change conformation to release the plug.

Computational modeling is a way of testing hypotheses regarding protein function in-silico. Compared to lab experiments or microscopy, computational testing provides a consistent and easily-manipulatable environment. Molecular dynamics is very useful for simulating molecular environments and introducing custom conditions [11]. Steered MD simulations with an applied force on the plug protein showed the required conformational change in DotG allowing for the release of the plug and other effector proteins.

To validate that our simulation was an accurate representation of the molecular environment, we embedded the protein in a membrane. The membrane was created using a custom-built Python package [12] with a concentration of lipids that resembled the actual lipid conformation in *L. pneumophila*. We repeated the steered MD with the membrane environment and observed similar simulation results. We concluded that our sans-membrane simulations are still accurate while being less computationally expensive.

Our conclusion, based on the results of our simulation, is that it is energetically and physically feasible that the plug protein could be pushed through the opening of DotG and out of the cell because of an external force applied by a build-up of effector proteins in the T4SS channel. The force-time series of both simulations is shown in Figure 5 and Figure 7.

**Figure 5:**
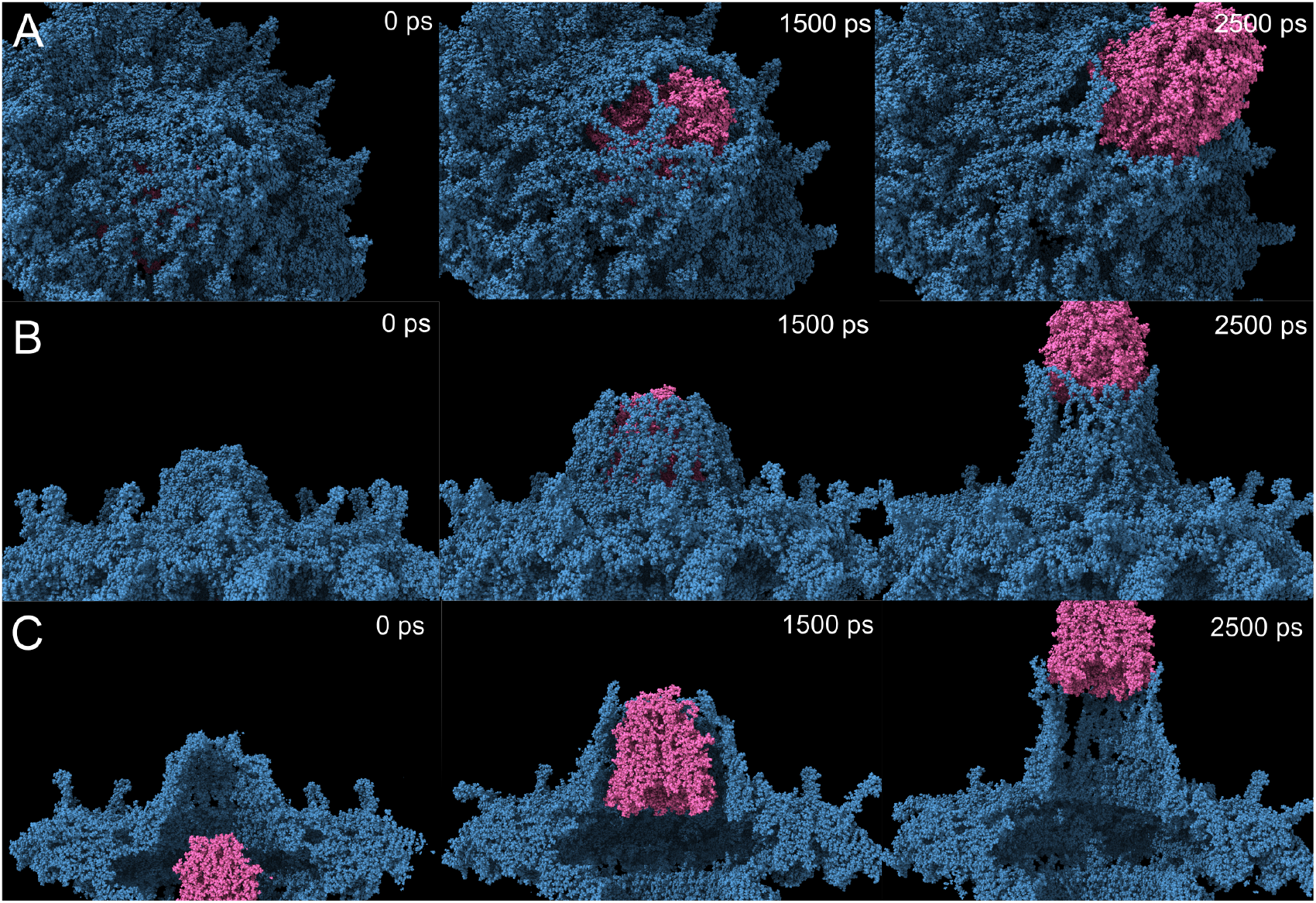
DotG Flexibility During Plug Release. (A) DotG flexibility at 0 ps, 1500 ps, and 2500 ps into the simulation of the release of the plug. Snapshots were taken from the simulation trajectory pushign the plug at 10 pm/ps. (B) Side view. (C) Cross section view.

**Figure 6:**
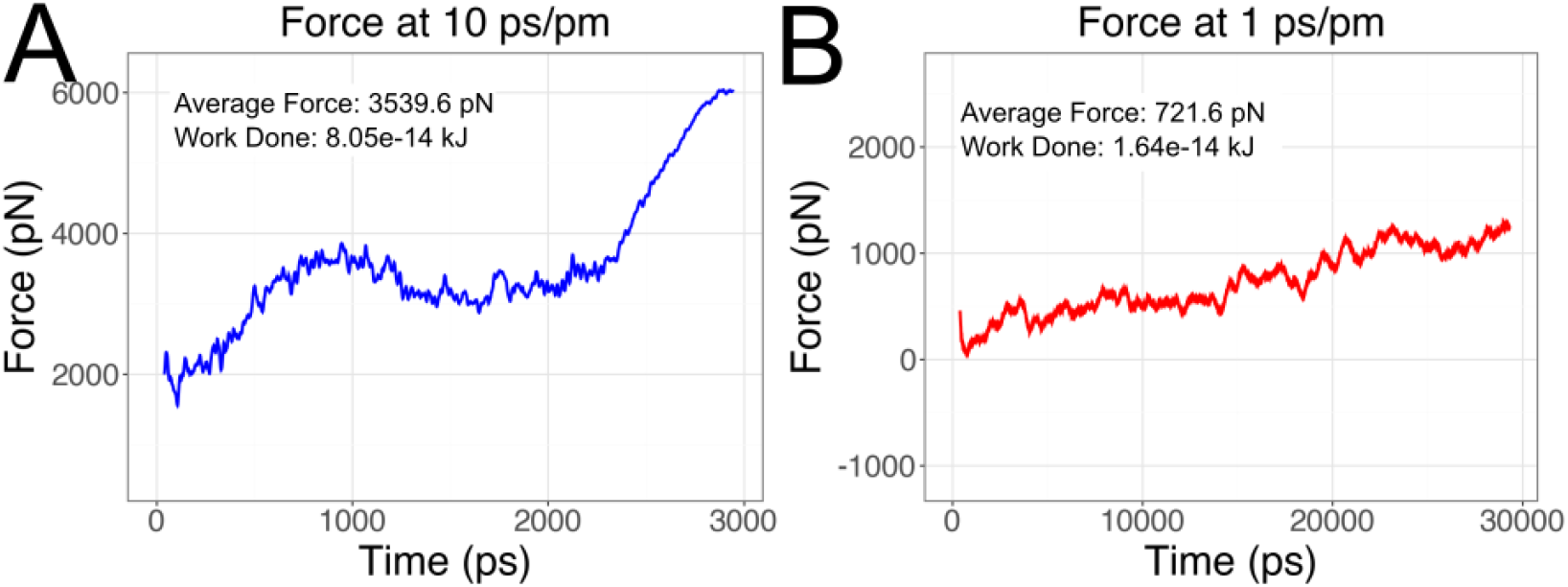
Required Secretion Forces. (A) The force required throughout the simulation to move the plug at a constant rate of 10 pm/ps. The average value of the force was calculated and thus the work was calculated given that the plug travels ∼ 220 Åthroughout the course of the simulation. (B) The force required to move the plug at a constant velocity of 1 pm/ps. Note that simulation had to be run for ten times longer so that the plug could still be fully secreted.

**Figure 7:**
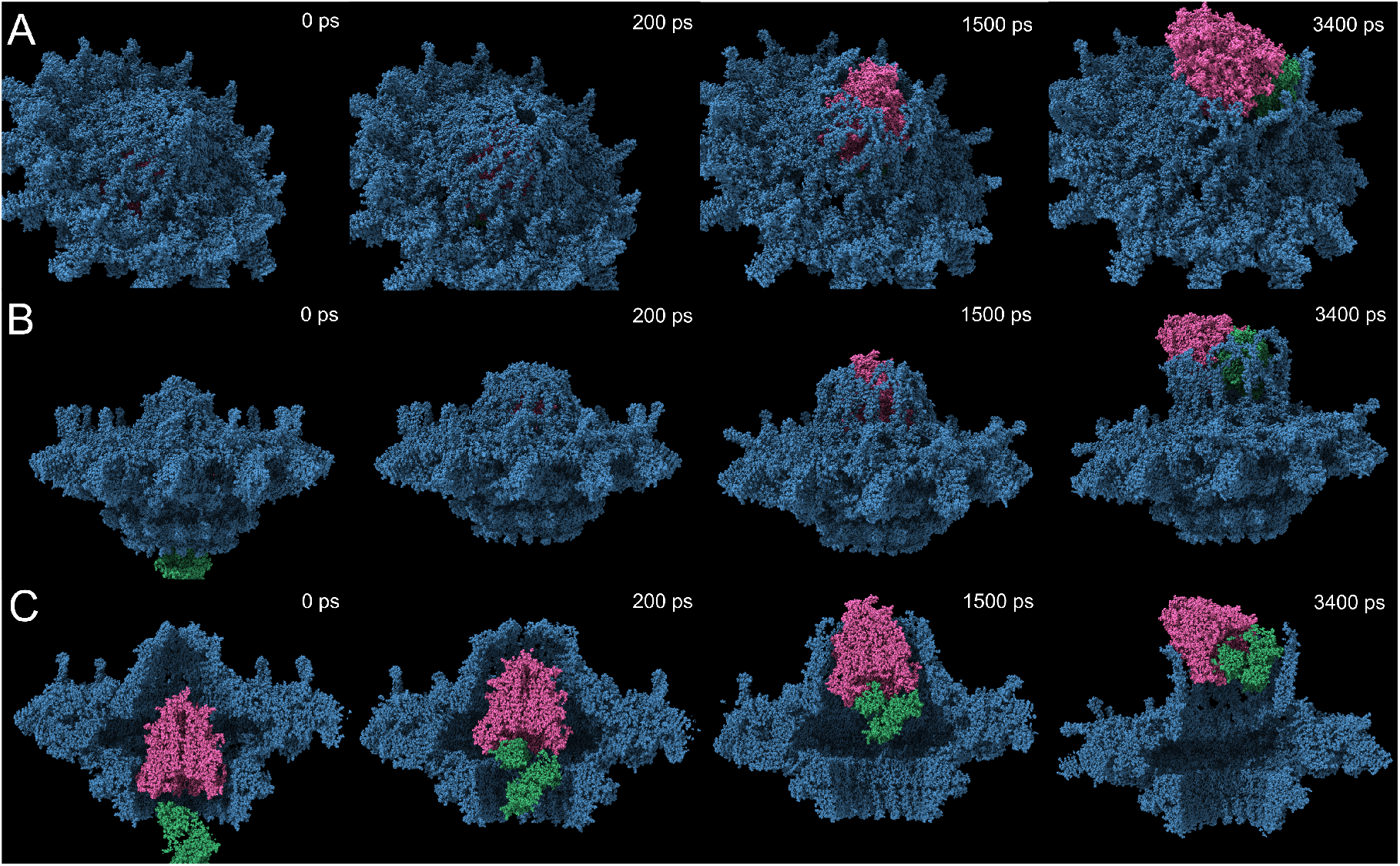
Effector and Plug Release at 5 pm/ps. (A) OMC simulation snapshots at 0 ps (beginning confirmation), 200 ps (contact between the effector protein and the plug), 1500 ps (plug begins to emerge from OMC), and 3400 ps (end of simulation). (B) Side view. (C) Cross section view.

## 3 Methods

### 3.1 Non-membrane Systems

Due to the large size of the T4SS complex, simulations focused on the region proximal to the outer membrane, including the outer membrane complex (OMC), periplasmic ring, and plug domain. This focus significantly reduced computational cost and allowed simulations to be conducted at accelerated pulling velocities.

Common imperfections in Protein Data Bank (PDB) files, such as missing heavy atoms and incomplete side chains, were addressed using OpenMM’s PDBFixer tool [13] and UCSF ChimeraX, which rebuilt side chains using most probable conformations according to the Dunbrack rotamer library [14].

Each system was placed in a periodic simulation box and solvated with SPC/E water [15]. The system’s net charge was neutralized using chloride counterions. These preprocessing steps were conducted using GROMACS utilities [16].

A cubic simulation box was generated around the protein complex, with a minimum clearance of 3 nm in each direction, resulting in a total volume of approximately 45 × 45 × 45 nm^3^. For simulations without the plug, the clearance was reduced to 2 nm to decrease system size. This box size ensured that the plug, once displaced, would not interact with its periodic image.

To evaluate the behavior of the system post-secretion, an extended simulation box of 38 × 38 × 105 nm^3^ was constructed to track plug motion over longer distances. However, as the additional space did not allow sufficient time for DotG to reset, these results were omitted from further analysis.

To assess whether the mechanical force required for plug secretion could plausibly arise from effector buildup, a known effector protein (Sidl) [17] was inserted below the plug. The force was applied to the effector protein’s center of mass instead of directly to the plug. The Sidl structure was inserted using ChimeraX. Sidl is a tRNA mimic that inhibits host protein synthesis via ribosome glycosylation [18].

### 3.2 Membrane Systems

A realistic outer membrane composition was approximated using a 5:3:2 ratio of POPE:POPC:POPG lipids, based on experimental data from *L. pneumophila* [19]. Standard membrane insertion tools were insufficient due to the system’s size and shape, so a custom Python package MembraneBuilder [12] was developed to generate tightly packed lipid bilayers. Lipids were initially constrained to high density and allowed to relax during energy minimization and equilibration, facilitating realistic interweaving. Membrane simulations were run analogously to the non-membrane simulations.

### 3.3 Molecular Dynamics Protocol

All molecular dynamics simulations were performed with GROMACS [16]. CHARMM27 was used for non-membrane simulations [20], and CHARMM36 for membrane systems, with additional lipid parameters sourced from the CHARMM-GUI Lipid Library [21, 22].

Each production simulation was run for 3400 ps with a time step of 2 fs. Long-range electrostatics were treated using the particle mesh Ewald (PME) method [23], with a cutoff of 12 Å and a switching function starting at 10 Å. Temperature was maintained at 300 K with the velocity-rescale thermostat, and pressure was controlled using the Parrinello–Rahman barostat [24].

Initial energy minimization was performed using the steepest descent algorithm for up to 20,000 steps, followed by NVT equilibration at 300 K for 50 ps.

### 3.4 Steered Molecular Dynamics (SMD)

To simulate plug secretion, we employed a steered molecular dynamics (SMD) [25] approach using a moving harmonic restraint applied between the center of mass (COM) of the plug and the rest of the complex. This was implemented using GROMACS pull code with ‘pull-coord1-type = umbrella’, but with a specified ‘pull-rate’ along the *z*-axis, consistent with non-equilibrium SMD.

Two pulling velocities were tested: 1 pm/ps and 10 pm/ps. These rates allowed us to compare force profiles and approximate the work done using numerical integration of the force-displacement curve. While this technique enables qualitative insight into the mechanical ejection of the plug, it is important to note that quantitative free energy estimates would require multiple independent trajectories. It is further important to emphasize the acceleration of the secretion, which would take 1,000-1,000,000 times longer in nature. Our simulations, limited to one replicate per pulling velocity, should therefore be interpreted with caution.

An analogous protocol was used for the effector simulation, with the harmonic pulling restraint applied to the COM of the effector protein rather than the plug.

### 3.5 Analysis

Trajectory visualization and inspection were performed using ChimeraX. The force applied during the pulling simulations was extracted from GROMACS-generated ‘.xvg’ files and analyzed using the `plotnine` and `ggplot` packages in Python [26]. Work was estimated by numerical integration of the force over the pulling distance. No error bars or confidence intervals are reported, as single trajectories are insufficient for statistical uncertainty estimation.

## 4 Results and Discussion

### 4.1 OMC Stability

Stationary simulations were run with and without the plug to determine whether the channel could remain stable without the presence of the plug. The ending conformations, at 3400 ps, from each simulation are shown in Figure 4. There was no notable difference in the system with and without the plug. The full simulation videos are available here: No Plug, With Plug.

### 4.2 Plug secretion through DotG

It is well known that effector proteins are transported by the T4SS into the host cell, but the mechanisms through which the proteins are secreted is poorly understood. Recent advances in cryo-electron tomography have led to the development of a complete molecular model of the protein system [2, 7, 8]. This model revealed the existence of a plug protein (IcmX), that sits in the transport channel. The lack of evidence of an extended pilus suggests that the effector proteins may be secreted through the opening of the T4SS channel (DotG), pushed through by some external force. This type of secretion would require a certain level of flexibility or change in structure of DotG to allow the effector proteins to pass through. We ran MD simulations to test the hypothesis that an external force on the plug could induce a conformational change in DotG and allow the plug and other effector proteins to pass through.

### 4.3 DotG Flexibility

Our primary simulation involved an induced force generated through umbrella pulling between the center of mass of the system and the plug to push it directly towards DotG. As the plug came in contact with DotG, the 16 chains in the DotG polymer flexed outwards and allowed for the secretion of the plug.

As can be seen in Figure 5, as the plug is pushed towards the opening of the secretion channel, the protein chains that form DotG flex outwards enough for the plug to pass through. A further observation is that the chains in the 16 member polymer DotG tend to group together while flexing. Full videos of the simulation can be seen here: Plug Secretion at 10 pm/ps.

### 4.4 Required Forces

In order to ensure this method is reasonable, the secretion simulation should require forces of a similar magnitude to similar processes in-vivo. The Type IV Pilus has been shown to extend at a speed near 0.5 *µ*m*/s* [27] and studies estimate the required force from these motors to exceed 100 *pN* [28]. Compared with our two simulations, this represents 7 and 6 orders of magnitude lower speed and only 1 or no orders of magnitude lower forces. Other secretion systems suggest varied force levels, such as the T3SS around 12-20 *pN* [29] and the T6SS reaching upwards of 500 *pN* [30]. Simulating the forces and speeds observed in-vivo would require a significantly larger computational and time cost. The force trend in our simulations (10 pm/ps to 1 pm/ps) shows an order of magnitude decrease. This suggests we would see realistic forces with realistic speeds.

The force required to pull the applied harmonic potential at a constant velocity was recorded throughout the simulation. This force was analyzed for two different secretion speeds, one at 10 pm/ps, and the other at 1 pm/ps. The average force was calculated and used to estimate the total work done given that the plug traveled ∼ 220 Å in the simulation. The force values were calculated at each 1 fs time step, and a moving average with a smoothing window (40 ps in the 10 pm/ps case, 400 ps in the 1 pm/ps case) was applied to reduce noise.

In both cases the force increased fairly consistently throughout the simuation. In the faster case, there was more variability in the force throughout the run. There is a particularly sharp increase in force necessary towards the end of the run, likely corresponding to the plug making its final push through DotG. This coincides with a steady increase in number of hydrogen bonds through the simulation, from *∼*20,500 to *∼*21,000.

In the slower case (1 pm/ps), the force required to push the plug through DotG has less variability, and there is a steady increase in the required force. The average force drops from 3539.6 pN to 721.6 pN, being reduced by a factor of ∼ 5 (4.9). The 5 times reduction in force despite having a 10 times reduction in speed, suggests that a slower speed promotes even more interaction and hydrogen bonding with the surrounding proteins.

A further reduction in speed would likely lead to a further reduction in the force required, approaching that of similar processes in-vivo (12-500 *pN*) [28], [29], [30].

### 4.5 Effector Protein Simulation

We hypothesized that this external force could reasonably come from the build-up of effector proteins pushed into the secretion channel by ATPases in the cytoplasmic section of the T4SS. We ran another simulation where the harmonic potential was placed on an effector protein located below the plug. The effector protein selected was Sidl, a protein that has been shown to mimic tRNA in the host cell, glycosylate ribosomes, and inhibit protein synthesis [18].

As can be seen in Figure 7, the effector protein simulation resembles the results from the standard simulation. The upward applied force caused Sidl to make contact with the bottom of the plug and push it upwards towards DotG. This interaction pushed both proteins out of the system in a similar way to what was initially observed with only the plug. The flexibility of DotG is once again critical to the secretion. The full simulation videos can be found here: Effector Plug Secretion 5 pm/ps.

Figure 8 shows the forces from the effector protein simulation and once again we see a reasonable average force given the secretion speed. This leads us to believe that the build-up of effector proteins in the channel can provide the required force to facilitate secretion. As before, the force values were smoothed with a window of 40 ps.

**Figure 8:**
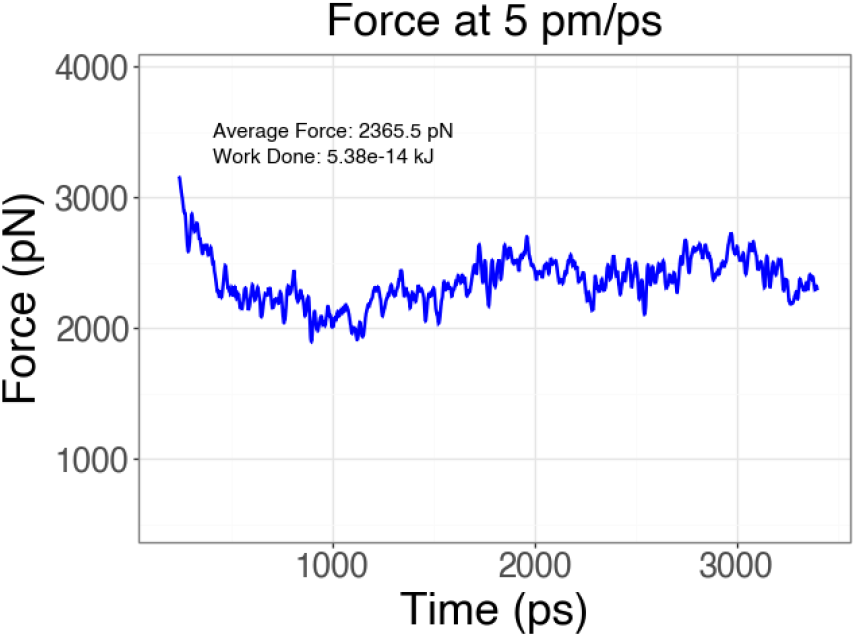
Force present during effector protein simulation. Note that the first 200 ps are removed from the graph to eliminate erroneously large forces at the beginning of the simulation. The applied harmonic potential was placed between the center of mass of the effector protein, and the center of mass of the rest of the system (excluding the plug). The effector protein began well below the rest of the system, and thus the harmonic potential to begin was extremely high.

### 4.6 Membrane System Results

To ensure our simulations were physiologically representative, we replicated the plug secretion simulation but with the system embedded in a membrane. We compared the trajectory of the secretion through the membrane with the sans-membrane secretion, and observed nearly identical reactions from the surrounding proteins–particularly the flexibility of DotG. Further, the average force throughout the membrane simulation (run at 10 pm/ps) was ∼ 3800 pN, compared to the ∼3500 pN from the sans-membrane simulation (also run at 10 pm/ps). While the membrane was not able to be equilibrated for an ideal length, we conclude that the sans-membrane environment is a valid model of the system and most further exploration excluded the membrane for computational convenience. The low equilibration time becomes evident in the gaps forming near the start of the membrane simulation. The full membrane system simulation can be viewed here: Membrane System Plug Secretion 10 pm/ps.

## 5 Conclusion

The simulations suggest that the Dot/Icm T4SS is capable of secreting effector proteins through the outer membrane complex, specifically DotG. It seems energetically feasible that the IcmX plug, which may be secreted first, can be pushed through the channel by the effector proteins moving through the channel. The forces required to push the plug through the channel suggest reasonable results given the secretion speed of the simulations. The flexibility of DotG is a key component in the secretion process, creating space for the plug to pass through.

These results provide more perspective on the secretion process of the Dot/Icm T4SS. The simulations provide a hypothetical look at the process of secretion and the involved forces. The results of this study can provide guidance in further understanding the function of the Dot/Icm T4SS. Future insights into the T4SS may provide a possible direction for developing new drugs to combat *Legionella pneumophila* infections. Drugs that inhibit DotG flexibility might theoretically prove effective at preventing effector protein secretion and thus render *Legionella pneumophila* incapable of evading host cell defenses and causing significant infection.

## Notes

https://www.youtube.com/@BYU_Biophysics_MD

### Competing Interest Statement

The authors have declared no competing interest.

